# Effects of Tempo, Dynamics, and String on Physical Exposure in Professional Violinists

**DOI:** 10.64898/2026.06.29.735269

**Authors:** Xuelong Fan, Svend Erik Mathiassen, Peter J. Johansson, Jennie A. Jackson, Teresia Nyman

**Author notes:** These authors contributed equally as co-senior authors (Jennie A. Jackson, Teresia Nyman).

## Abstract

1.

This study examined how tempo, dynamics, and string influence upper-extremity physical exposure in professional violinists and how exposure variability is distributed among musical characteristics, between-subject differences, and residual variability. Twelve violinists performed seven standardized scales while bilateral upper-arm and wrist kinematics and shoulder and forearm muscle activity were recorded. Linear mixed-effects models showed that faster tempo increased right upper-arm velocity and bilateral forearm activity while reducing right upper-arm and wrist ranges of motion. Louder dynamics increased bilateral forearm and right trapezius activity and right-wrist ranges of motion. Higher-posture strings increased right upper-arm elevation and right shoulder muscle activity. Variance analysis identified exposures predominantly related to musical characteristics, jointly related to musical characteristics and between-subject differences, predominantly related to between-subject differences, or mainly unexplained. These findings support future exposure prediction from musical characteristics and targeted prevention through repertoire-based workload management, structured recovery, and individualized technique-focused strategies.

**Highlights:**

- Tempo, dynamics, and string produced distinct upper-extremity exposures.
- Faster tempo and louder dynamics increased bilateral forearm muscle activity.
- Higher-posture strings increased right upper-arm elevation and shoulder activity.
- Distinct musical, between-subject, and residual variance patterns were identified.
- Findings support repertoire-based and individualized PRMSD prevention.

## 3. Introduction

Occupational musculoskeletal disorders (MSDs), often referred to as playing-related MSDs (PRMSDs) in musicians, are prevalent among classical musicians (Cruder et al., 2020, 2023; Leaver et al., 2011; Stanhope et al., 2022; Vastamäki et al., 2020). The most common pain locations reported by university level or professional musicians include the neck, shoulders, and lower back (Leaver et al., 2011). Professional orchestra musicians are 50% more likely to develop back pain and 160% more likely to develop shoulder pain than the general workforce (Vastamäki et al., 2020). For musicians, occupational exposure includes time spent practicing alone, rehearsing with other musicians, recording music, and performing in front of an audience. While the overall practicing load has been shown to be dependent on experience, for violinists, it generally includes 4–6 hours per day and 17–28 hours per week (Chi et al., 2020)

Among classical musicians, upper string players, including violinists and violists, have a higher prevalence of PRMSDs than musicians playing other instruments (Kok et al., 2018; Leaver et al., 2011; Nyman et al., 2007; Paarup et al., 2011). Playing an upper string instrument involves elevated and unsupported arms performing asymmetric tasks requiring static loading, repetitive movements, and precise motor control. The non-bowing arm (the left arm) stabilizes the instrument and performs repeated shoulder rotations and precise finger movements to depress the strings in exact locations while producing rhythmic oscillation of the fingertip to produce vibrato. The bowing arm (the right arm) performs repetitive movements involving the shoulder, elbow, wrist, and fingers to draw the bow across the strings with precise motor control required to produce any specific type of sound (ex. quiet vs loud, and slow vs fast). Furthermore, the two upper extremities must also continuously coordinate postural support, fine motor control, and force regulation to arrive at the desired sound. This bilateral coordination may increase physical exposure across both sides of the upper body, particularly during technically demanding passages of music.

Upper-extremity exposures are likely influenced by the characteristics of the music played, including the speed at which notes are played (tempo), the volume of the sound generated (dynamics), and the string on which the notes are played. Previous studies have examined the effects of tempo and dynamics on shoulder and forearm muscle activity; however, these studies have included musicians with different proficiency levels, and their results may not reflect the exposure patterns of highly skilled professional musicians (Mann et al., 2021; Mann, Paarup, et al., 2023). Less is known about how these musical characteristics affect kinematic physical exposures, including upper-arm and wrist postures and movement velocities, exposures that have previously been associated with neck and shoulder musculoskeletal disorders in other types of work (Nordander et al., 2016). Tempo is expected to influence movement velocity, particularly in the bowing arm and in the fingers of the non-bowing hand. Dynamics are expected to influence the amount of force required to produce sound, with playing louder, i.e., fortissimo (ff), requiring greater bowing force than playing softer, i.e., pianissimo (pp). The string played is expected to affect both posture and muscle activity of both the bowing and non-bowing arms, as illustrated in **Figure 1**. The placement order of the four strings of a violin, E, A, D, and G, dictate the playing posture bilaterally, where playing on the E-string (the string with the highest pitch) requires the lowest right shoulder elevation and the largest external rotation of the left shoulder, and playing on the G-string (the string with the lowest pitch) requires the highest shoulder elevation and the smallest external rotation of the left shoulder). Together, these biomechanical considerations suggest that musical characteristics may shape violinists’ intrinsic physical loads, including upper extremity posture, movement velocity, and muscle activity (Rensing et al., 2018). Identifying how tempo, dynamics, and string choice affect these playing-related exposures is therefore important for characterizing workload in violinists, understanding PRMSD mechanisms, and developing targeted prevention strategies.

**Figure 1.**
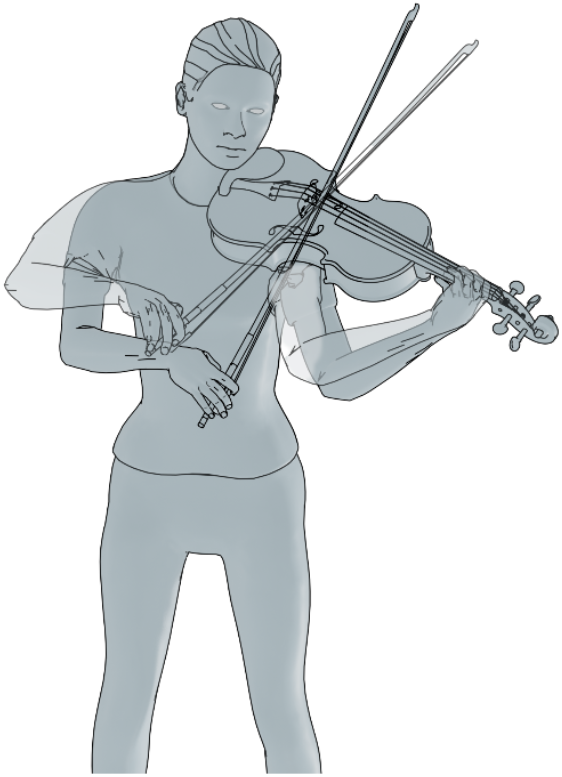
Illustration showing the change in upper extremity postures in a violinist playing on different strings, with the darker grey position showing postures while playing on the E-string (the string with the highest pitch, the lowest right arm elevation, and the largest external rotation of the left shoulder) and the light grey position showing the postures while playing on the G-string (the string with the lowest pitch, the highest arm elevation and the smallest external rotation of the left shoulder).

In addition to musical characteristics, individual playing technique has been shown to affect the biomechanical exposures of professional violinists, where a high intra-subject repeatability has been reported for violinists performing the same piece of music twice (Fjellman-Wiklund et al., 2004; Mann, Paarup, et al., 2023; Visentin et al., 2015). However, little is known about the relative extent to which exposure variability *between* violinists is attributable to musical characteristics versus individual factors. Such information is needed as a first step towards being able to predict exposures while playing complex pieces of music, and thus developing a basis for, e.g., structuring rehearsal plans and choice of repertoire.

Hence, the primary aim of this study was to investigate the effects of three musical characteristics, i.e., tempo, dynamics, and string, on upper extremity posture, movement velocity, and muscle activity in professional violinists. The secondary aim was to investigate the proportion of the variance in each physical exposure that was attributed to musical characteristics relative to that residing in individual playing technique.

## 4. Materials and methods

### 4.1. Participants

Twelve professional violinists, employed in a Swedish symphony orchestra or working freelance and who were not currently on sick leave, participated in the study. All participants were informed about the study protocol, provided written informed consent, and completed a screening regarding musculoskeletal pain experienced during the past seven days (yes/no) and the past twelve months (never/seldom/sometimes/often/very often) (Jonker et al., 2015). The study was approved by the Regional Ethical Review Board in Uppsala (Dnr 2018/412, 2018/412-1) and the Swedish Ethical Review Authority (Dnr 2019-04617).

### 4.2. Experimental protocol

Each violinist participated in a laboratory experiment, during which muscle activity and kinematics were recorded in the shoulders, upper arms, forearms, and wrists. Playing performances were simultaneously video recorded for reference.

The study followed an experimental crossover design. During the experiment, participants performed seven one-octave scales (eight ascending and eight descending notes) at 120 beats per minute with a time signature of 4/4. The seven scales systematically varied in three musical characteristics: tempo (2 levels: whole notes [30 notes per minute] and sixteenth notes [480 notes per minute]), dynamics (2 levels: pianissimo [pp] and fortissimo [ff]), and the strings in which the scales were played, operationalized using scale key as a proxy (3 levels: E major, A major, and G major).

In this study, the four violin strings were grouped as “higher-posture” and “lower-posture” strings, based on the right-arm, or bowing-arm, posture required to play them, rather than on their pitch (**Figure 1**). The E and A strings were therefore considered lower-posture strings, whereas the D and G strings were considered higher-posture strings. To complete a one-octave scale, the notes can be played either across two strings with the same left-hand position or on one string with a shift in left-hand position. The A-major scale was played on the A and E strings with the same left-hand position and was used as the lower-posture reference. The G-major scale was played on the G and D strings with the same left-hand position and was used as the higher-posture comparison. Thus, the comparison between the G- and A-major scales was treated as the primary contrast for assessing differences between higher- and lower-posture strings. The E-major scale was played solely on the E string but required a shift in left-hand position on the neck of the violin. It was therefore included as an additional lower-posture condition. Comparisons between the G- and E-major scales were used as additional higher-versus lower-posture contrasts, whereas comparisons between the E- and A-major scales were treated as secondary contrasts to examine differences related to left-hand position and playing exposure within lower-posture strings. (**Table 1**).

**Table 1.**
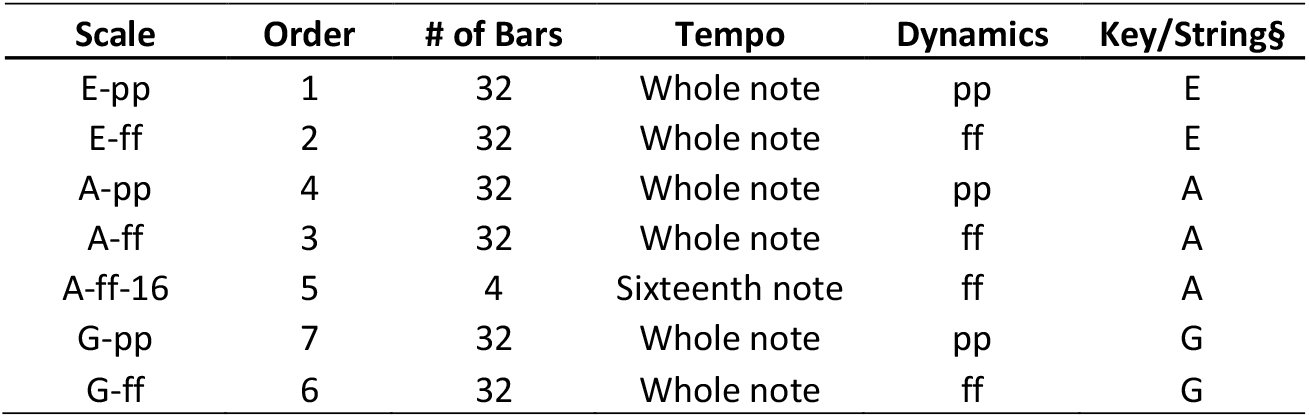
Descriptions of the seven scale tasks performed by the participants, defined by combinations of tempo, dynamics, and key/string. All scales were one-octave played with eight ascending and eight descending notes, at 120 beats per minute and a 4/4 time signature, with one note per bow stroke. Tempo was defined by note value: whole notes corresponded to 30 notes per minute, and sixteenth notes corresponded to 480 notes per minute. For dynamics, pp denotes pianissimo, and ff denotes fortissimo. All scales started with playing the open string, i.e., the E major scale started on an open E-string. § The scale key was used as a proxy for the strings played during each scale: The E major scale was played only on the E string, the A major scale on the A and E strings, and the G major scale on the G and D strings.

Additionally, a metronome was used throughout to ensure the speed was consistent. Participants were asked to play the scales as if performing them for an audience, using their best vibrato and producing their best sound. To ensure scales were played in the same manner across all participants, the fingering was provided for each scale. All scales were played with smooth movements and with the bow constantly on the string (called legato bowing). Participants were seated on a chair without a backrest. Music sheet was placed on a stand positioned at a self-selected comfortable height and distance, and page turning was not required. Participants were given sufficient time to feel warmed up and to then rest before beginning the experiment.

### 4.3. Measurement of kinematics

Kinematic measurements of the upper arms were obtained using two accelerometers (AX3, Axivity Ltd., Newcastle, UK) recording at 25 Hz (Fan et al., 2021). The sensors were positioned beneath the deltoid muscles on both upper arms.

Bilateral wrist kinematics were recorded using two biaxial goniometers (Biometrics Ltd., Cwmfelinfach, Gwent, UK) at a sampling frequency of 1000 Hz. The goniometer was placed with one block over the third metacarpal bone, and the other block positioned at the midline between the two forearm bones, as presented in Simonsen et al. (Simonsen et al., 2018).

Upper-arm elevation was defined as inclination angles relative to a neutral vertical reference posture performed by each participant while sitting with a lateral lean until the upper arm hung vertically while holding a 2 kg dumbbell (Bernmark & Wiktorin, 2002). The positive direction in the elevation/depression movement for the upper arm was elevation. The reference posture for the wrist (0 degrees flexion and deviation) was defined by having the participants place their hands flat on a horizontal surface, with the hand adjusted so that a straight line was formed between the third metacarpal bone and the forearm, as presented by Nordander et al (Nordander et al., 2016). The maximal flexion/extension range of motion of the wrist was determined by having the participant perform maximal wrist flexion and extension three times while seated with their elbows supported on a table and the forearms positioned horizontally. Maximal ulnar and radial deviation range of motion were determined for each participant through three voluntary range of motion trials performed with the palm flat on a horizontal surface. The positive direction in the flexion/extension movement for the wrist was defined as flexion, while that in the deviation movement was defined as radial deviation.

### 4.4. Measurement of muscle activity

Muscle activity of the upper trapezius, deltoid, and forearm muscles was monitored bilaterally using surface electromyography (sEMG). Self-adhesive bipolar Ag/AgCl electrodes (N-00-S/25, Ambu A/S, Copenhagen, Denmark) were applied with an inter-electrode distance of 2 cm at each location. Before electrode placement, the skin was shaved and cleaned with alcohol to reduce impedance. Electrodes were secured using conductive gel and adhesive fixation to maintain contact during movement. The signals were transmitted via actively shielded cables to a digital data logger (Oldenzaal, The Netherlands), sampled at 1000 Hz per channel using a 24-bit A/D converter.

For the upper trapezius, electrodes were positioned 2 cm lateral to the midpoint between the C7 spinous process and the acromion (Mathiassen et al., 1995). For the deltoid, electrodes were placed according to the SENIAM (Surface EMG for the Non-Invasive Assessment of Muscles) guidelines for the middle/lateral deltoid at the midpoint between the acromion and the deltoid tuberosity along the muscle belly (Hermie J. Hermens et al., n.d.). Forearm muscle activity was recorded using a through-forearm placement of sEMG electrodes (Fan et al., 2026; Takala & Toivonen, 2013). This electrode placement was selected to capture the combined activity from both forearm extensors and flexors, thereby enhancing the accuracy of forearm workload estimation. Specifically, electrodes were placed just lateral to the midpoint of the belly of the extensor digitorum communis muscle and just lateral to the belly of the flexor digitorum superficialis muscle (Takala & Toivonen, 2013).

Submaximal reference voluntary electrical activations (RVE) trials were recorded during the reference voluntary contractions (RVCs). Reference contractions for the upper trapezius and deltoid were performed with participants standing in a neutral posture, elevating both arms to 90 degrees (Mathiassen et al., 1995). For the forearm muscles, submaximal reference contractions were obtained one side at a time with the forearm supported on a table from elbow to wrist, the hand extended over the edge of the table, and a 1 kg weight hung from a soft strap that was positioned across the third proximal interphalangeal joint of the fingers on the hand. Each reference contraction was repeated three times for each participant. All RVC trials were 15 s in duration, interspersed by 30 s of rest (Åkesson et al., 1997; G.-Å. Hansson et al., 2000).

### 4.5. Data processing

The EMG signals were processed through the following processing pipeline. First, the raw signals were digitally bandpass-filtered between 20 and 400 Hz and then passed through a 50 Hz comb notch filter to remove power-line interference. The root mean square (RMS) values of the filtered signals were computed using 125-ms epochs and subsequently corrected for noise by power subtraction. The resulting RMS values were normalized to the corresponding RVE level, calculated as the mean RMS value from the three RVC trials, and expressed as %RVE (Mathiassen et al., 1995). The 50th percentile (P50) was calculated for each muscle site and for each scale. (Fan et al., 2024; G.-Å. Hansson et al., 2000).

Accelerometer data were low-pass filtered at 5 Hz using a Blackman window filter and subsequently converted into elevation angles for the upper arms, with positive values indicating elevation. The 50th percentile was calculated as a summary metric for elevation. Ranges of motion were computed as the difference between the 90th and 10th percentiles (P90 – P10) of the angle distributions. Angular velocities were obtained as the first derivatives of the corresponding angular signals, as described in previous studies (Fan et al., 2021; G. Hansson et al., 2001), and 50th percentile values were calculated even for velocity.

Goniometer signals were resampled to 20 Hz, low-pass filtered at 5 Hz using a Blackman window filter, and calibrated by subtracting the wrist angle measured in the neutral reference posture (G.-Å. Hansson et al., 1996), yielding deviation and flexion angles, which were used to calculate the range of motion and angular velocity. Consequently, data processing yielded one numeric value for each scale and physical exposure measure for each participant, and these values were subsequently used in the statistical analyses.

### 4.6. Statistical analysis

All analyses were conducted in MATLAB R2023a (The MathWorks, Inc., Natick, MA, USA).

Descriptive results of measured physical exposures were summarized and presented in box plots and tables, and pairwise differences within each musical characteristic were summarized as median differences with first and third quartiles.

Associations between musical characteristics and physical exposures were evaluated using linear mixed-effects (LME) models in a repeated-measures design accounting for both the effects of the musical characteristics (fixed effects) and the between-subject variability (random effects). Thus, in the LME models, the physical exposure measures were treated as outcomes, while musical characteristics (tempo, dynamics, and string) were set as fixed effects, and subjects were a random intercept. The significance level was set at α = 0.05.

Model construction followed a forward, likelihood-based stepwise procedure with strong heredity for interaction terms. The base model, *measure*_*ij*_ = *β*_0_ + *u*_*i*_ + *ϵ*_*ij*_, was fitted using maximum likelihood (ML) to allow comparisons between models with different fixed-effect structures and was treated as the compact model. Interaction terms were considered between fixed effects only after inclusion of the corresponding main effects, and were restricted to the dynamics × string interaction, in accordance with the study design. At each step, a single candidate term from the set of the remaining candidate terms was added independently to the compact model to form a full model; the resulting set of models was compared using a likelihood-ratio test (LRT) under ML. The candidate with the smallest p-value was retained if p < α (primary criterion). When multiple candidates showed comparable evidence near the threshold, changes in the Bayesian Information Criterion (ΔBIC) were used as a secondary criterion, favoring greater reductions in BIC. The process was repeated until the process reached a point where no additional candidate met the inclusion threshold α or no candidates were left. The maximal model that can be reached in theory was as follows (based on the current study design, E.1):

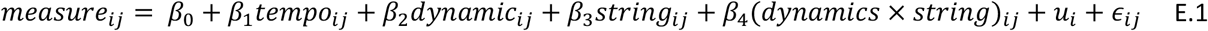

where *i* indexes participants and *j* indexes observations, *β*_0_ represents the fixed intercept, and *β*_1−4_ represent the fixed-effect coefficients for tempo, dynamics, string, and the interaction between dynamics and string. *u*_*i*_ denotes the participant-specific random intercept, and *ϵ*_*ij*_ denotes the residual error.

For each outcome, residuals from the base model were screened for homoscedasticity (primary) and normality (secondary). Homoscedasticity was assessed using the Breusch–Pagan test and a median-based Levene test across fitted-value quartiles (Breusch & Pagan, 1979; Brown & Forsythe, 1974). Homoscedasticity was concluded only when both tests were non-significant (p ≥ α). Normality was evaluated using the Shapiro–Wilk test (Shapiro & Wilk, 1965). When heteroscedasticity was detected, a monotonic Box–Cox transformation with an optimally estimated λ was applied, and model selection was repeated on the transformed scale (Box & Cox, 1964). Otherwise, analyses were performed on the original scale, given the relative robustness of LME models to moderate deviations from normality (Blanca et al., 2017).

After fixed-effect selection, the final linear mixed-effects model was refitted using restricted maximum likelihood (REML) to obtain less biased variance-component estimates and final parameter estimates for the selected fixed effects. Post hoc pairwise comparisons were then conducted to evaluate categorical fixed effects of the three musical characteristics. For two-level factors (i.e., tempo and dynamics), the model coefficient directly represented the adjusted mean difference. For factors with more than two levels (i.e., string), pairwise contrasts were computed using linear combinations of fixed-effect estimates derived from the fitted model. P-values were adjusted for multiple comparisons using Bonferroni correction.

When models were fitted using a Box–Cox transformation to address distributional assumptions, statistical inference was performed on the transformed scale. For interpretability, model-predicted values were back-transformed to the original scale using the estimated transformation parameter (λ), and model-performance parameters, such as root mean square errors (RMSEs), were computed on the original scale using the inverse Box–Cox function.

To quantify the relative contributions of musical characteristics, between-subject differences, and residual variability, variance metrics were calculated following established methods, including marginal R^2^ (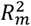; fixed effects), conditional R^2^ (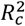; fixed and random effects), and the intraclass correlation coefficient (ICC) (Nakagawa & Schielzeth, 2013). Additionally, 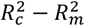 and 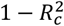 were calculated to quantify the proportions of variance attributable to between-subject differences and residual variability. Together, 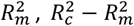, and 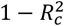 partitioned the total variance into components attributable to musical characteristics, between-subject differences, and residual variability.

Larger 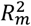 values indicate a stronger contribution of the investigated musical characteristics; larger 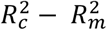 values indicate greater between-subject variability captured by the random intercept; and larger 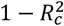 values indicate greater residual variability. The ICC represents between-subject variance relative to between-subject plus residual variance, with larger values indicating greater between-subject variability relative to residual variability. Although these measures are mathematically related and contain overlapping information, 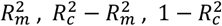, and the ICC were examined jointly to facilitate identification of recurring variance patterns across exposures. The equations for these variance measures are provided in Eqs. 2–6:

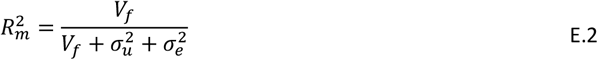

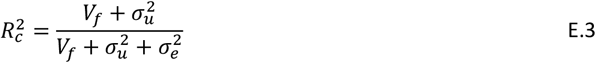

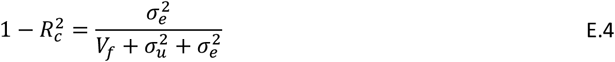

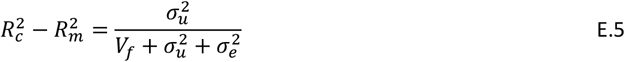

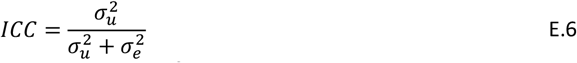

where *V*_*f*_ is the variance of the fixed-effects predictions, 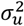 is the total variance of the random effects, and 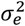 is the residual variance estimated from the model.

## 5. Results

On average, participants were 49 years old, 171 cm tall, and weighed 70 kg. More than half of the participants were female (58%), and most participants were right-handed (91%). All held the violin in their left hand and held the bow in their right hand. More than half (58%) of the participants had had pain in at least one body part in the last 7 days, and half reported having had pain often or very often in at least one upper extremity area part in the last 12 months. Nonetheless, all participants stated that they were not on sick leave and could play normally as they intended (**Table 2**). All 12 participants completed all scales.

**Table 2.**
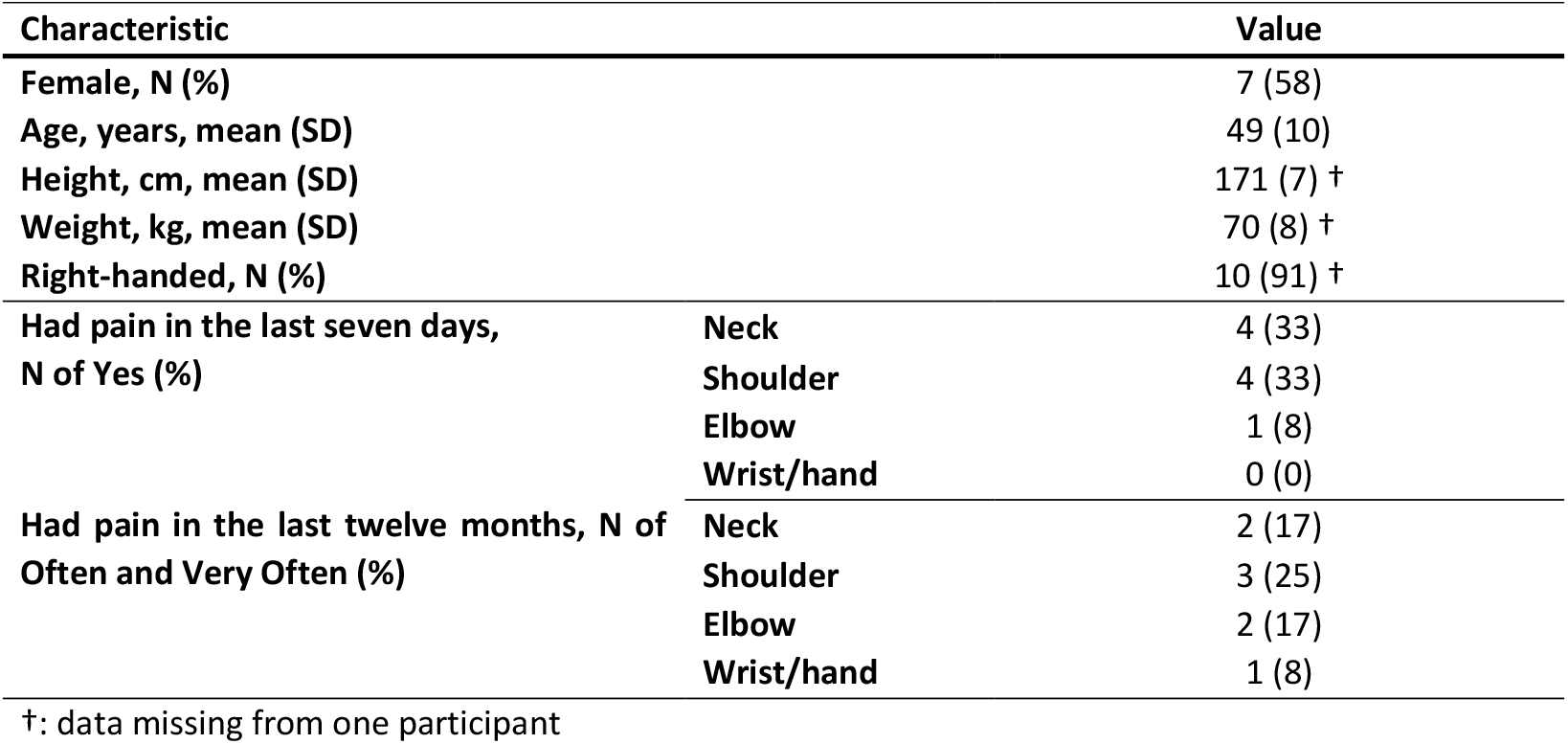
The demographic characteristics of the participants.

| Characteristic | Value |
| --- | --- |
| Female, N (%) | 7 (58) |
| Age, years, mean (SD) | 49 (10) |
| Height, cm, mean (SD) | 171 (7) † |
| Weight, kg, mean (SD) | 70 (8) † |
| Right-handed, N (%) | 10 (91) † |
| Had pain in the last seven days, N of Yes (%) |  |
| Neck | 4 (33) |
| Shoulder | 4 (33) |
| Elbow | 1 (8) |
| Wrist/hand | 0 (0) |
| Had pain in the last twelve months, N of Often and Very Often (%) |  |
| Neck | 2 (17) |
| Shoulder | 3 (25) |
| Elbow | 2 (17) |
| Wrist/hand | 1 (8) |

†: data missing from one participant

All three musical characteristics, i.e., tempo, dynamics, and string, affected kinematic and muscle activity physical exposure metrics, but in different ways. For all exposure metrics, descriptive results, including distribution of values, median values, and first and third quartile values, are shown in **Figure 2** and **Figure 3**, and a complete set of numerical results is shown in **Table S1. Tables 3–5** show within-subject differences for each musical characteristic, and **Table 6** shows the results of the statistical tests of these differences and variance-related metrics. These tables are described in more detail below.

**Table 3.** Median differences in physical exposure between different music tempi. L/R indicates the left or right side; Q1 and Q3 denote the first and third quartiles; and the reference (Ref) is the whole note.

| Tempo: Sixteenth note vs (Ref) Whole note, Difference, Median [Q1, Q3] |  |  |  |  |  |  |  |  |  |
| --- | --- | --- | --- | --- | --- | --- | --- | --- | --- |
| Section | Posture |  |  | Posture (°) | Range of motion (°) | Velocity (°/s) | Muscle | Muscle activity (%RVE) |  |
|  |  |  |  | P50 | P90 - P10 | P50 |  | P50 |  |
|  |  |  |  | Difference | Difference | Difference |  | Difference |  |
| Shoulder/<br>Upper arm | Upper arm | Elevation | L | -5 [-6, -1] | -2 [-4, 0] | 6 [3, 13] | Deltoid | L | 2 [-6, 6] |
|  |  |  | R | -9 [-10, -6] | -11 [-16, -4] | 26 [23, 40] |  | R | 14 [8, 18] |
|  |  |  |  |  |  |  | Trapezius | L | 1 [-8, 12] |
|  |  |  |  |  |  |  |  | R | -27 [-39, -10] |
| Forearm/<br>Wrist | Wrist | Deviation | L | -4 [-6, -3] | -2 [-4, 0] | 5 [3, 7] | Through* | L | 40 [-2, 46] |
|  |  |  | R | -2 [-4, 2] | -12 [-18, -6] | 6 [3, 11] |  | R | 48 [15, 66] |
|  |  | Flexion | L | -6 [-11, -4] | -6 [-8, -3] | 4 [0, 6] |  |  |  |
|  |  |  | R | 1 [-3, 5] | -34 [-36, -22] | -3 [-17, 1] |  |  |  |

\* Forearm muscle activity measured by the through-forearm EMG placement

**Figure 2.**
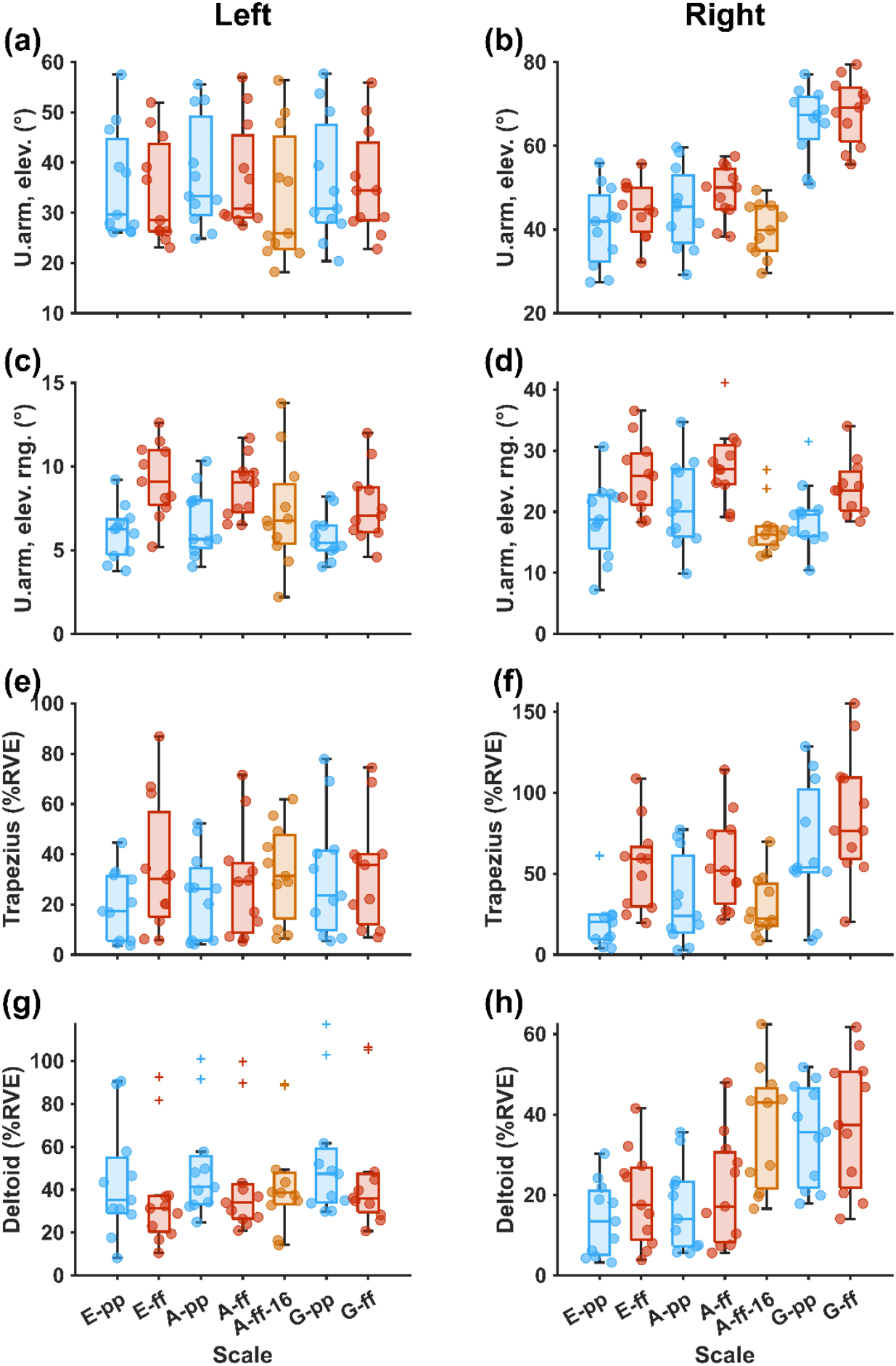
Boxplots of the median (P50) exposures of the upper arms (u.arm) and shoulder muscles on both the right and left sides under different musical characteristics. Elev. stands for elevation, and rng. stands for range. The central line within each box indicates the median, while the box edges represent the 1st and 3rd quartiles, and the whiskers denote the minimum and maximum values. Round markers indicate individual data points, and symbols marked “+” denote Tukey outliers. Blue markers and bars represent scales played whole notes at pianissimo; red markers and bars represent whole notes at fortissimo; and yellow markers and bars represent scales played with sixteenth notes.

**Figure 3.**
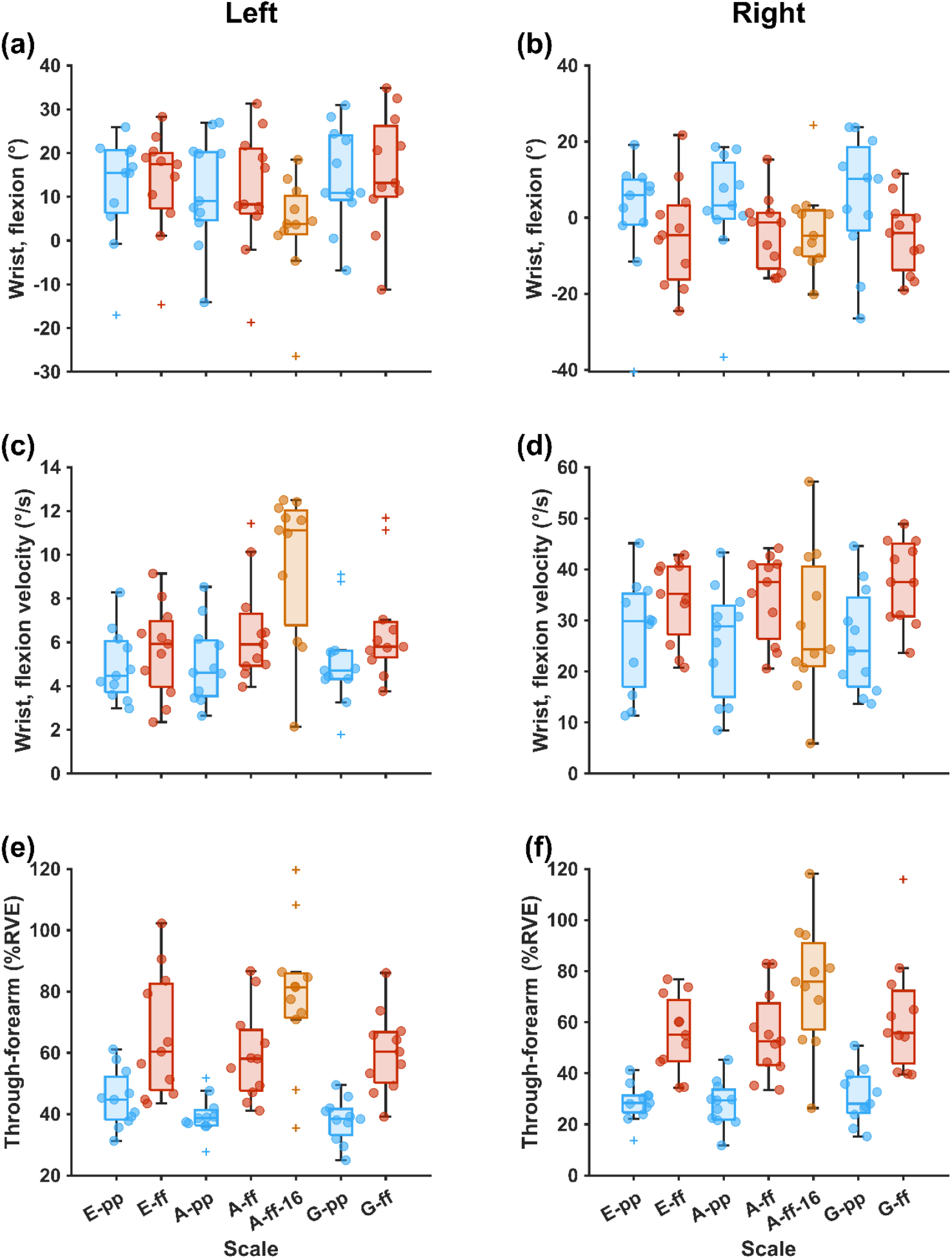
Boxplots of the median (P50) exposures of the wrists and lower arms on both the right and left sides under different musical characteristics. The central line within each box indicates the median, while the box edges represent the 1st and 3rd quartiles, and the whiskers denote the minimum and maximum values. Round markers indicate individual data points, and “+” symbols denote Tukey outliers. Blue markers and bars represent scales played whole notes at pianissimo; red markers and bars represent whole notes at fortissimo; and yellow markers and bars represent scales played with sixteenth notes.

From the modelled data, no interaction between dynamics and string was found (all p > 0.7).

### 5.1. Effects of tempo

Changing the tempo from whole notes to sixteenth notes (from slow to fast) produced clear changes in kinematic and muscle activity, especially on the right side (**Figure 2, Figure 3, Table 3, Table 6**). The median velocity of the right upper-arm elevation increased by 26°/s ([Q1, Q3], [23, 40]; p < 0.001), the median elevation angle decreased by 9° ([−10, −6]; p < 0.001), and the median range of elevation (P90–P10) decreased by 11° ([−16, −4]; p < 0.001). These kinematic changes were accompanied by altered shoulder muscle activity, including a 14 %RVE ([8, 18]; p < 0.001) increase in median right deltoid activity and a 27 %RVE ([−39, −10]; p < 0.001) decrease in median right trapezius activity.

In the right forearm, the median range of right-wrist deviation and flexion decreased by 12° ([−18, −6]; p < 0.001) and 34° ([−36, −22]; p < 0.001) (**Table 3, Table 6**) and the median forearm muscle activity in the bowing (right) arm increased by 48 %RVE ([15, 66]; p < 0.001).

The effects of tempo on the kinematics and muscle activity of the left side were either non-significant or small in magnitude (**Table 3, Table 6**) with the exception of the left forearm muscle activity, which increased by 40 %RVE ([-2, 46]; p = 0.001) when playing at the faster tempo.

Overall, kinematic and muscle activity changes in both the shoulder and forearm regions occurred when playing at a faster tempo and were predominantly evident in the bowing (right) arm, except for forearm muscle activity, which increased bilaterally.

### 5.2. Effects of dynamics

Dynamics also affected both kinematic measures and muscle activity (**Figure 2, Figure 3, Table 4, Table 6, Table S1**). On the right side, louder dynamics increased median trapezius activity by 25%RVE ([Q1, Q3] [8, 37]; p < 0.001) and forearm muscle activity by 24%RVE ([12, 39]; p < 0.001). Louder dynamics also increased the range of right wrist deviation by 10° ([5, 14]; p < 0.001) and the range of right wrist flexion by 14° ([4, 21]; p < 0.001). Other right-sided kinematic measures also changed significantly, although the magnitudes were relatively small. Dynamics also affected the left side; however, the only sizeable change when playing louder (*ff*) compared to softer (*pp*) was observed in forearm muscle activity, which increased by 23%RVE ([11, 40]; p < 0.001) (**Table 4, Table 6**). Other changes in physical exposures were minor in magnitude or non-significant.

**Table 4.** Median differences in physical exposure between different music dynamics. L/R indicates the left or right side; Q1 and Q3 denote the first and third quartiles; and the reference (Ref) is pp.

| Dynamics: ff vs (Ref) pp, Difference, Median [Q1, Q3] |  |  |  |  |  |  |  |  |  |
| --- | --- | --- | --- | --- | --- | --- | --- | --- | --- |
| Section | Posture |  |  | Posture (°) | Range of motion (°) | Velocity (°/s) | Muscle | Muscle activity (%RVE) |  |
|  |  |  |  | P50 | P90 - P10 | P50 |  | P50 |  |
|  |  |  |  | Difference | Difference | Difference |  | Difference |  |
| Shoulder/<br>Upper arm | Upper arm | Elevation | L | -1 [-2, 0] | 2 [1, 4] | 2 [1, 3] | Deltoid | L | -8 [-11, -1] |
|  |  |  | R | 3 [0, 7] | 6 [3, 8] | 6 [3, 7] |  | R | 3 [0, 8] |
|  |  |  |  |  |  |  | Trapezius | L | 2 [0, 7] |
|  |  |  |  |  |  |  |  | R | 25 [8, 37] |
| Forearm/<br>Wrist | Wrist | Deviation | L | 1 [-1, 1] | 0 [0, 3] | 0 [0, 1] | Through* | L | 23 [11, 40] |
|  |  |  | R | 4 [-1, 7] | 10 [5, 14] | 5 [3, 8] |  | R | 24 [12, 39] |
|  |  | Flexion | L | 1 [-1, 2] | 2 [0, 3] | 1 [1, 2] |  |  |  |
|  |  |  | R | -9 [-14, -3] | 14 [4, 21] | 8 [2, 14] |  |  |  |

\* Forearm muscle activity measured by the through-forearm EMG placement

Overall, dynamics predominantly affected right shoulder muscle activity, forearm muscle activity on both sides, and the range of motion of the right wrist.

### 5.3. Effects of string

As described in the Methods, string effects were evaluated using scale key as a proxy, with the G-versus A-major comparison serving as the primary contrast between higher- and lower-posture strings. Comparisons involving the E-major scale were used as additional contrasts to assess whether similar patterns were observed across lower-posture conditions and whether left-hand position influenced the results. Overall, string-related differences were most evident on the right side, whereas left-hand-position-related differences were more limited and mainly observed on the left side (**Figure 2, Figure 3, Table 5, Table 6, Table S1**).

**Table 5.**
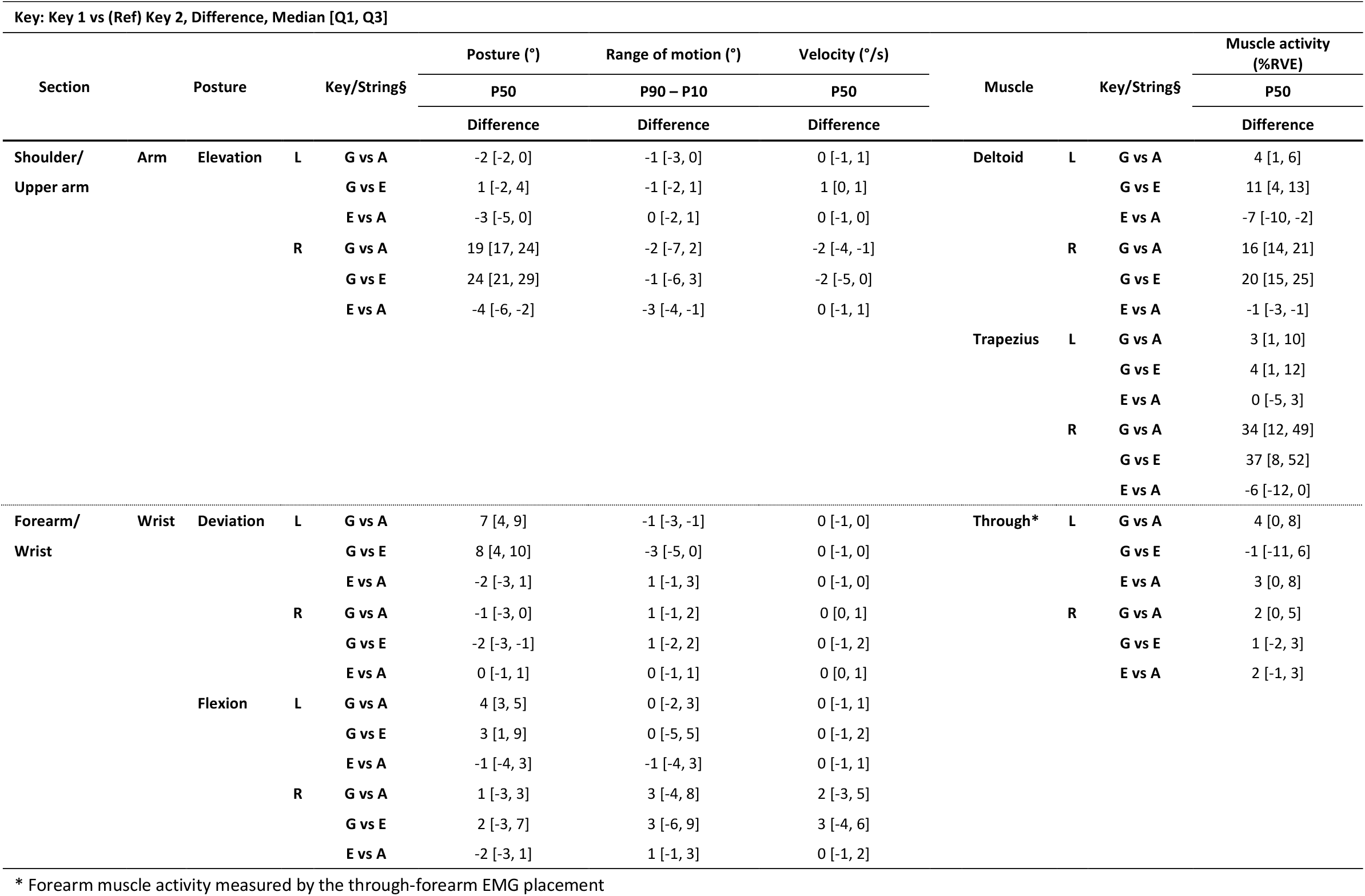
Median differences in physical exposure between different strings proxied by the scales played in different major keys. L/R indicates the left or right side; Q1 and Q3 denote the first and third quartiles; and the reference (Ref) is the key after “vs”. § The scale key was used as a proxy for the strings played during each scale.

The largest right-sided changes were observed when the G-major scale, which involved higher-posture strings, was compared with the A- and E-major scales, which involved lower-posture strings (**Table 5, Table 6**). Compared with the A-major scale, the G-major scale increased right upper-arm elevation by 19° ([Q1, Q3] [17, 24]; p < 0.001). Compared with the E-major scale, the G-major scale increased right upper-arm elevation by 24° ([21, 29]; p < 0.001). In contrast, no significant difference was observed between the A- and E-major scales, suggesting that right upper-arm elevation was mainly associated with differences in strings used during the scale rather than left-hand position.

**Table 6.** Effects of the three musical characteristics and variances explained by the models. RMSE denotes the root-mean-square error; 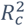 and 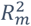 represent the conditional and marginal R^2^, respectively; and ICC indicates the intraclass correlation coefficient. P-values are Bonferroni-adjusted for pairwise contrasts. P<0.05 is considered significant. § The scale key was used as a proxy for the strings played during each scale.

| Variable | Body section | Direction | Side | Tempo | Dynamics | Key/String§ | | | | RMSE | $R_m^2$ | $1 - R_c^2$ | $R_c^2 - R_m^2$ | ICC |
| --- | --- | --- | --- | --- | --- | --- | --- | --- | --- | --- | --- | --- | --- | --- |
|  |  |  |  | 16th vs whole | ff vs pp | Overall | G vs A | G vs E | E vs A |  |  |  |  |  |
| Posture<br>(°) | Wrist | Deviation | L | <0.001 | - | <0.001 | <0.001 | <0.001 | >0.999 | 3.1 | 0.19 | 0.15 | 0.66 | 0.82 |
|  |  |  | R | - | <0.001 | - | - | - | - | 3.3 | 0.10 | 0.30 | 0.60 | 0.67 |
|  |  | Flexion | L | <0.001 | - | 0.009 | 0.010 | 0.133 | 0.980 | 3.8 | 0.08 | 0.10 | 0.82 | 0.89 |
|  |  |  | R | - | <0.001 | - | - | - | - | 6.5† | 0.07 | 0.24 | 0.68 | 0.74 |
|  | Upper arm | Elevation | L | <0.001 | - | 0.002 | 0.098 | 0.456 | 0.002 | 2.0 | 0.01 | 0.04 | 0.95 | 0.96 |
|  |  |  | R | <0.001 | <0.001 | <0.001 | <0.001 | <0.001 | <0.001 | 3.2 | 0.65 | 0.07 | 0.28 | 0.80 |
| Range of motion, (P90 - P10)<br>(°) | Wrist | Deviation | L | 0.034 | - | 0.007 | 0.087 | 0.009 | >0.999 | 4.3† | 0.07 | 0.39 | 0.53 | 0.57 |
|  |  |  | R | <0.001 | <0.001 | - | - | - | - | 4.0 | 0.39 | 0.27 | 0.34 | 0.55 |
|  |  | Flexion | L | <0.001 | - | - | - | - | - | 4.4† | 0.12 | 0.63 | 0.26 | 0.29 |
|  |  |  | R | <0.001 | <0.001 | - | - | - | - | 8.4 | 0.44 | 0.28 | 0.29 | 0.51 |
|  | Upper arm | Elevation | L | 0.032 | <0.001 | - | - | - | - | 1.8† | 0.20 | 0.63 | 0.17 | 0.21 |
|  |  |  | R | <0.001 | <0.001 | - | - | - | - | 4.5 | 0.25 | 0.52 | 0.22 | 0.30 |
| Velocity<br>(°/s) | Wrist | Deviation | L | <0.001 | 0.002 | - | - | - | - | 1.3† | 0.37 | 0.42 | 0.21 | 0.34 |
|  |  |  | R | <0.001 | <0.001 | - | - | - | - | 4.9† | 0.42 | 0.37 | 0.22 | 0.37 |
|  |  | Flexion | L | <0.001 | <0.001 | - | - | - | - | 1.4† | 0.21 | 0.31 | 0.48 | 0.61 |
|  |  |  | R | 0.007 | <0.001 | - | - | - | - | 6.2 | 0.15 | 0.36 | 0.49 | 0.58 |
|  | Upper arm | Elevation | L | <0.001 | <0.001 | - | - | - | - | 3.0† | 0.47 | 0.14 | 0.39 | 0.73 |
|  |  |  | R | <0.001 | <0.001 | 0.004 | 0.024 | 0.008 | >0.999 | 8.2† | 0.69 | 0.19 | 0.12 | 0.40 |
| Muscle activity<br>(%RVE) | Through* |  | L | 0.001 | <0.001 | - | - | - | - | 13.2† | 0.49 | 0.40 | 0.12 | 0.23 |
|  |  |  | R | <0.001 | <0.001 | - | - | - | - | 13.3† | 0.57 | 0.32 | 0.11 | 0.26 |
|  | Deltoid |  | L | - | <0.001 | <0.001 | 0.090 | <0.001 | 0.002 | 5.9 | 0.03 | 0.06 | 0.91 | 0.94 |
|  |  |  | R | <0.001 | 0.004 | <0.001 | <0.001 | <0.001 | 0.310 | 5.8† | 0.34 | 0.12 | 0.54 | 0.82 |
|  | Trapezius |  | L | - | 0.015 | - | - | - | - | 12.0 | 0.03 | 0.36 | 0.62 | 0.63 |
|  |  |  | R | <0.001 | <0.001 | <0.001 | <0.001 | <0.001 | 0.578 | 15.8† | 0.33 | 0.23 | 0.44 | 0.66 |

\* Forearm muscle activity measured by the through-forearm EMG placement
† Estimated on the Box-Cox-transformed scale and back-transformed to the original outcome scale.

A similar pattern was observed for right shoulder muscle activity (**Table 5, Table 6**). Right trapezius activity increased by 34%RVE when comparing the G-major scale with the A-major scale ([12, 49]; p < 0.001) and by 37%RVE when comparing the G-major scale with the E-major scale ([8, 52]; p < 0.001), with no significant difference between the A- and E-major scales. Right deltoid activity increased by 16%RVE from the A-major scale to the G-major scale ([14, 21]; p < 0.001) and by 20%RVE from the E-major scale to the G-major scale ([15, 25]; p < 0.001), again with no significant difference between the A- and E-major scales. Notably, the magnitude of the increase was consistently larger when comparing the G-major scale with the E-major scale than with the A-major scale. Other right-sided physical exposures were either non-significant or small in magnitude.

On the left side, changes were mainly observed in wrist deviation and deltoid activity (**Table 5, Table 6**). Similar to the right-sided pattern, left wrist radial deviation increased when comparing the G-major scale with the A-major scale by 7° ([4, 9]; p < 0.001) and with the E-major scale by 8° ([4, 10]; p < 0.001), while no significant difference was observed between the A- and E-major scales.

In contrast, left deltoid activity showed a different pattern (**Table 5, Table 6**). It increased by 11%RVE when comparing the E-major scale with the G-major scale ([4, 13]; p < 0.001), whereas the difference between the A- and G-major scales was not significant. Moreover, when comparing the two lower-posture conditions, the E-major scale was associated with a 7%RVE decrease in left deltoid activity compared with the A-major scale ([-10, -2]; p = 0.002). This suggests that left deltoid activity may be influenced more by left-hand position than by the strings used during the scale. Other left-sided exposures were either non-significant or small in magnitude.

Overall, differences between strings played in each scale were mainly reflected in right upper-arm elevation and right shoulder muscle activity, with moderate changes in left-wrist deviation. In contrast, differences in left-hand positions on similar lower-posture strings appeared to affect mainly left deltoid activity, with limited or no influence on right-sided exposures.

### 5.4. Variance analysis and model performance

Variance analysis identified four distinct patterns across the physical exposure variables (**Table 6, Figure S1**). The four patterns represented exposures predominantly related to musical characteristics, jointly related to musical characteristics and between-subject differences, predominantly related to between-subject differences, or characterized by substantial unexplained variability, with little variation attributable to either the examined musical characteristics or between-subject differences.

The first pattern showed the strongest dependence on musical characteristics. It was characterized by moderate-to-large 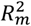 values (0.37–0.69), low-to-moderate 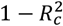 values (0.18–0.42), small 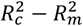 values (0.11–0.22), and low-to-moderate ICCs (0.23–0.40). This pattern was most evident for right upper-arm elevation velocity, which had 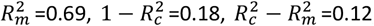, and ICC = 0.40. Bilateral forearm muscle activity showed a similar pattern, with 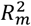 values of 0.48–0.57, 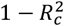 values of 0.32– 0.40, 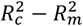 values of 0.11–0.12, and ICCs of 0.23–0.26. Bilateral wrist-deviation velocity also belonged to this pattern. These exposures were therefore influenced substantially by musical characteristics, while between-subject differences accounted for only a small proportion of the total variance. In most cases, the unexplained residual variance exceeded the variance attributable to between-subject differences.

The second pattern reflected substantial contributions from both musical characteristics and between-subject differences. It combined moderate-to-large 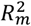 values (0.33–0.65), relatively small 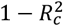 values (0.07–0.28), and 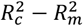 values (0.28–0.54) with moderate-to-high ICCs (0.51–0.81). Right upper-arm elevation posture showed this pattern, with 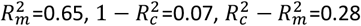, and ICC = 0.80. A similar pattern was observed for left upper-arm elevation velocity, right-wrist deviation and flexion ranges of motion, and right deltoid and trapezius activity. For these exposures, musical characteristics and between-subject differences together explained most of the variance, leaving comparatively little residual variability.

The third and most common pattern was dominated by between-subject differences. It was characterized by small 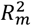 values (0.01–0.21), low-to-moderate 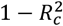 values (0.04–0.40), moderate-to-large 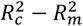 values (0.48–0.95), and moderate-to-high ICCs (0.57–0.96). For example, left upper-arm elevation posture had 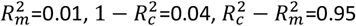, and ICC = 0.96. Left deltoid activity showed a similar pattern ( 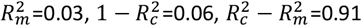, ICC = 0.94). This pattern was also observed for left trapezius activity, bilateral wrist postures, bilateral wrist-flexion velocity, and left wrist-deviation range of motion. For these exposures, between-subject differences accounted for the largest proportion of variance, while musical characteristics made a comparatively limited contribution.

The fourth pattern was characterized by substantial unexplained variability. It comprised left wrist-flexion range of motion and bilateral upper-arm elevation range of motion, which had relatively small 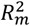 values (0.12–0.26), large 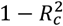 values (0.52–0.63), small 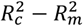 values (0.17–0.26), and low ICCs (0.21–0.30). Thus, neither musical characteristics nor between-subject differences explained a large proportion of the variance in these three exposures; instead, most variability remained unexplained by the models.

RMSEs were generally small relative to the magnitude of the corresponding outcomes, particularly for the kinematic measures, indicating acceptable average prediction error at the group level (**Table 6, Table S1**). Overall, the variance analysis distinguished exposures predominantly related to musical characteristics, jointly related to musical characteristics and between-subject differences, predominantly related to between-subject differences, or largely dependent on residual factors not captured by the models.

## 6. Discussion

This study examined how three musical characteristics—tempo, dynamics, and string—were associated with upper-extremity kinematics and muscle activity in professional violinists. All three characteristics affected physical exposure, but in distinct ways.

Faster tempo primarily affected the right, or bowing, side, increasing right upper-arm velocity and forearm muscle activity while reducing right upper-arm and wrist ranges of motion. Louder dynamics mainly increased muscle activity in both forearms and the right upper trapezius and increased right-wrist ranges of motion. Playing on higher-posture strings mainly affected the right side, increasing right upper-arm elevation and right shoulder muscle activity, whereas effects related to left-hand position were limited and were observed mainly in left deltoid activity.

Variance analysis further showed that the sources of variability differed across exposures. Right upper-arm elevation velocity, bilateral forearm muscle activity, and wrist-deviation velocity were predominantly related to musical characteristics. Right upper-arm elevation posture, left upper-arm elevation velocity, right-wrist ranges of motion, and right shoulder muscle activity reflected contributions from both musical characteristics and between-subject differences. In contrast, many wrist postures, wrist-flexion velocities, and left shoulder exposures were predominantly related to between-subject differences. Finally, left wrist-flexion range of motion and bilateral upper-arm elevation range of motion were characterized mainly by residual variability not explained by the investigated musical characteristics or between-subject differences.

### 6.1. Effects of music characteristics on exposure

Each musical characteristic influenced physical exposure in distinct patterns, suggesting different underlying mechanisms.

Tempo primarily affected right-sided kinematics and muscle activity. Faster tempo increased right upper-arm velocity and muscle activity in the right deltoid and both forearms, while reducing the ranges of motion of the right upper arm and wrist. These findings are partly consistent with previous studies showing increased physical demand at higher tempi (Mann, Paarup, et al., 2023; Rensing et al., 2018; Reynolds et al., 2014), but extend them by demonstrating concurrent reductions in ranges of motion in both the right wrist and upper arm. This combination suggests a movement strategy characterized by faster but more constrained motion, potentially limiting unnecessary joint movement and supporting rapid and precise bowing. A notable contrast to this combination was the decrease in right trapezius activity despite increased right deltoid and forearm activity. Earlier studies reported either no change or increased right trapezius activity at higher tempi (Mann, Paarup, et al., 2023; Reynolds et al., 2014). Because previous studies included mixed-skill populations, whereas the present study involved only professional violinists, the discrepancy may reflect skill- and experience-dependent motor strategies. Professional violinists may redistribute muscular effort within the shoulder complex during faster playing, reducing trapezius activity while maintaining bowing performance.

Dynamics mainly modulated muscle activity, particularly in both forearms and the right upper trapezius. This pattern partly aligns with previous findings of increased right-forearm activity and left-finger force at stronger dynamics (Kinoshita & Obata, 2009; Mann, Paarup, et al., 2023; Rensing et al., 2018), but the present results also showed clearer bilateral forearm engagement. Unlike tempo, stronger dynamics increased the ranges and velocities of right-wrist flexion and deviation. This suggests that louder playing may rely on wider and faster distal joint ranges of motion to help forearm and shoulder muscles generate bow force and sound intensity, whereas changes in upper-arm posture were comparatively small.

Although string use was assessed using scales played in different keys, the results suggest that the strings used during the scale were the main driver of right-sided exposure differences. The G-major scale was played on higher-posture strings, namely the G and D strings, whereas the A-major scale was played on lower-posture strings, namely the A and E strings. The E-major scale represented another lower-posture condition, as it was played only on the E string but with two different left-hand positions. As string position became higher, right-sided shoulder exposure showed a graded increase, with the G-major scale producing the greatest right upper-arm elevation and right deltoid and upper-trapezius activity, followed by the A-major and then the E-major scales. This is consistent with previous reports that shoulder movement increases when playing shifts from the E string toward the G string (Shan & Visentin, 2003; Wolf et al., 2019). In contrast, differences between the A- and E-major scales had limited effects on right-sided exposures, suggesting that neither left-hand position alone nor the shift between the A and E strings was sufficient to substantially alter right-arm demands.

On the left side, the string-related pattern differed. Left wrist deviation changed when the G-major scale was compared with the other keys, showing a pattern broadly consistent with the right-sided response. However, left deltoid activity differed between the A- and E-major scales and increased significantly only between the G- and E-major scales. This suggests that left deltoid activity may be more sensitive to left-hand position or instrument-stabilizing demands than to the strings used alone.

Thus, across musical characteristics, both similarities and contrasts emerged. Tempo and dynamics both increased forearm muscle activity, indicating that both demanded force generation and control. However, they differed in their kinematic patterns: an increase in tempo was associated with reduced joint ranges of motion, whereas louder dynamics increased wrist ranges of motion. In contrast, string effects were primarily proximal, affecting upper-arm elevation and shoulder muscle activity, and reflected positional requirements rather than dynamic or temporal demands. Together, these findings suggest that the investigated characteristics influenced physical exposure through largely distinct mechanisms.

### 6.2. Variance explained by musical characteristics, between-subject differences, and residual variability

A key finding of this study was that physical exposures differed not only in their responses to the investigated musical characteristics, but also in how their total variance was distributed among musical characteristics, between-subject differences, and residual variability. Joint consideration of 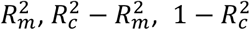, and the ICC identified four distinct patterns: exposures predominantly related to musical characteristics, exposures jointly related to musical characteristics and between-subject differences, exposures predominantly related to between-subject differences, and exposures characterized mainly by residual variability.

The first pattern represented the strongest dependence on musical characteristics and was observed for right upper-arm elevation velocity, bilateral forearm muscle activity, and bilateral wrist-deviation velocity. For these exposures, tempo, dynamics, or string accounted for a substantial proportion of the total variance, whereas between-subject differences contributed comparatively little. This suggests that these physical exposures are strongly shaped and constrained by the demands imposed by the musical characteristics, allowing relatively little variation in average exposure levels between violinists. Although musical characteristics explained more variance than between-subject differences, a substantial proportion of the variance remained unexplained for bilateral forearm muscle activity and bilateral wrist-deviation velocity. Therefore, these exposures cannot be considered fully predictable from the investigated musical characteristics alone, and additional factors not included in the present study likely contribute to their variability. The importance of this residual variability may differ across exposures. For example, wrist-deviation velocity, particularly on the left side, showed relatively limited variation across the investigated scales, suggesting that part of the unexplained variance may reflect measurement noise or short-term fluctuations with limited practical importance. In contrast, for bilateral forearm muscle activity, the residual component may reflect a combination of measurement-related variability and unmeasured aspects of performance. Previous studies have reported low-to-moderate variability in forearm EMG during simple and standardized manual tasks, indicating that some variability is inherent to EMG measurements themselves (Fan et al., 2026). In addition, subtle differences in sound quality, artistic expression, and other performance-related nuances requiring fine motor control of the forearms and hands may contribute to variability that was not captured by the present models and therefore appears as residual variance.

The second pattern reflected substantial contributions from both musical characteristics and between-subject differences. It was observed for right upper-arm elevation posture, left upper-arm elevation velocity, right-wrist deviation and flexion ranges of motion, and right deltoid and trapezius activity. Musical characteristics and between-subject differences together explained most of the variance in these exposures, leaving relatively little residual variability. Right upper-arm elevation posture contrasted particularly clearly with right upper-arm elevation velocity: both were strongly related to musical characteristics, but posture also showed substantial between-subject variability relative to its small residual component. Thus, the musical characteristics shifted these exposures systematically, while average exposure levels also differed between violinists. Such differences may reflect technique, anthropometry, habitual posture, instrument setup, or motor strategy, although the random-intercept models cannot determine the contribution of each factor.

The third and most common pattern was dominated by between-subject differences and included left upper-arm elevation posture, left deltoid and trapezius activity, bilateral wrist postures, bilateral wrist-flexion velocity, and left wrist-deviation range of motion. For these exposures, musical characteristics explained comparatively little variance, whereas differences between violinists accounted for the largest component. This suggests that their average levels may depend more strongly on individual posture, technique, anthropometry, instrument setup, or motor strategy than on the investigated musical characteristics. This interpretation is consistent with previous findings of substantial between-subject differences in trapezius activity during violin playing (Fjellman-Wiklund et al., 2004). The same study reported relatively low within-subject variability during repeated performances, suggesting that muscle-activity levels may remain comparatively consistent within individual musicians even when similar material is performed repeatedly. However, because the present models included random intercepts rather than random slopes, they describe differences in average exposure levels between violinists and do not establish that violinists responded differently to changes in musical characteristics.

The fourth pattern was characterized mainly by residual variability and comprised three exposures: left wrist-flexion range of motion and bilateral upper-arm elevation range of motion. Neither musical characteristics nor between-subject differences explained a large proportion of their variance. Similar to the first pattern, the unexplained variability exceeded the variability attributable to between-subject differences. In contrast to the exposures in the first pattern, however, these ranges of motion showed relatively small changes across the investigated scales, indicating that the examined musical characteristics had limited influence on them. Their variability may instead reflect short-term movement adjustments, certain technique (e.g., use of vibrato in the left hand), measurement variability, interactions not represented in the models, or other factors not captured in the present study, rather than systematic effects of musical characteristics or stable differences between violinists.

Together, these patterns show that physical exposures during violin performance differ in whether they are predominantly related to musical characteristics, jointly related to musical characteristics and between-subject differences, predominantly related to between-subject differences, or largely unexplained by the present models. This distinction has practical implications. Exposures strongly related to musical characteristics may be more predictable from repertoire and more amenable to workload management based on the duration and sequencing of demanding passages. Exposures with substantial between-subject contributions may require individualized attention to technique, posture, instrument setup, and motor strategy. For exposures dominated by residual variability, the unexplained variance may partly reflect factors not included in the present models, such as sound production, i.e., use of vibrato, artistic expression, and other performance-related nuances requiring fine motor control that were not measured in this study. Further studies are needed to identify additional explanatory factors and to determine the extent to which these performance-related factors contribute to the observed variability before targeted interventions are developed.

### 6.3. Physical exposures and risk of developing musculoskeletal disorders

The magnitude of a physical exposure is meaningful for PRMSD risk only when considered together with its temporal profile, including how long, how often, and in what sequence the exposure occurs. Many risk and exposure assessment studies focus on the whole-day exposure (6–8 hours) in occupations with relatively homogenous work tasks. For example, proposed action levels are exposure thresholds above which preventive measures should be considered to reduce the risk of work-related musculoskeletal disorders (Arvidsson et al., 2021). These thresholds are based on prolonged measurements, often covering an approximately 8-hour working day, whereas the present study examined short standardized scale tasks. However, scales with similar musical characteristics are recurring components of violinists’ daily practice, rehearsal, and performance. The observed task-specific exposures can therefore inform approximate exposure scenarios for days dominated by music with similar characteristics, although they should not be interpreted as directly equivalent to full-day occupational measurements.

Within this context, several exposures reached magnitudes comparable to those reported in physically demanding occupations. Median right upper-arm elevation involving lower-posture strings was generally approximately 30°–50° and increased to around 70° during the G-major scale involving higher-posture strings. These values are comparable to or larger than those reported in occupations such as hairdressing (29° at the 50^th^ percentile) and retail shelf stocking (55° at the 90^th^ percentile) (Forsman et al., 2017; Nordander et al., 2016). High arm elevation has been associated with an increased risk of shoulder disorders (Van Der Molen et al., 2017). Specifically, median right upper-arm elevation exceeded the proposed action level of 30° was considered as a risk factor for developing MSDs, which was lower than the right-arm elevation observed in most scales (Arvidsson et al., 2021). This indicates that elevated right-arm posture represents a substantial baseline exposure in violin playing, while music involving higher-posture strings may further increase the accumulated shoulder load.

Right trapezius activity also varied with musical characteristics. Playing louder increased trapezius activity, particularly when combined with higher-posture strings, whereas activity during playing softer was lower. Direct comparison with occupational action levels can be difficult because the present study used RVE normalization and task-specific exposure measures, whereas the proposed action levels are based on prolonged occupational measurements and MVE-normalized muscle activity. Nevertheless, the observed pattern indicates that louder dynamics and higher-posture strings may add substantially to shoulder muscle load, and therefore the risk of PRMSDs around neck and shoulder, when such passages are repeated or sustained over longer periods.

Taken together, the potential risk of PRMSDs in violinists is likely determined by the combination of consistently elevated baseline exposures and additional increases associated with specific musical characteristics, accumulated across practice, rehearsal, and performance. Because different musical characteristics produced distinct exposure patterns, some loads may be mitigated through repertoire variation during practicing and rehearsal, for example by alternating passages with different dynamics or passages involving higher- and lower-posture strings. However, the musical characteristics required within a given piece are often intrinsic to the repertoire and cannot be modified directly, especially during performance. Their temporal profile nevertheless provides an important degree of freedom for intervention. Demanding passages may be distributed across practice sessions, alternated with passages involving lower exposures, limited in uninterrupted duration, and separated by structured recovery periods. Thus, even when the magnitude of exposure associated with particular musical characteristics cannot be reduced, workload-management strategies can target its duration, frequency, sequencing, and recovery pattern.

### 6.4. Practical implications

The present findings have potential implications for occupational workload management based on repertoire selection and therefore strategies for preventing PRMSDsat an organizational level.

First, the results demonstrate that different musical characteristics impose distinct biomechanical demands, meaning that physical exposure can vary substantially depending on what music piece is being played, even when practice duration remains constant. This is particularly relevant in professional orchestras, where rehearsal schedules are typically fixed despite variations in repertoire (Nyman et al., 2007). Introducing variation in practice and rehearsal schemes based on musical characteristics, such as alternating between passages with different dynamics or between music played predominantly on lower and higher strings, may help distribute mechanical load across rehearsal periods and reduce localized fatigue and disorder risk.

Second, the diverse variance patterns across physical exposures indicate that different risk-mitigation strategies may be required. Exposures predominantly related to musical characteristics, such as right upper-arm elevation velocity and bilateral forearm muscle activity, may be partly managed through repertoire variation and scheduling. Identifying these characteristic-dependent exposures also provides a basis for estimating physical exposure from musical characteristics and may ultimately support exposure prediction directly from sheet music. In contrast, right upper-arm elevation posture remained consistently high across conditions, although its variance reflected contributions from both musical characteristics and between-subject differences. Repertoire selection alone is therefore unlikely to reduce this exposure substantially while conventional violin-playing techniques are maintained. A combination of repertoire-based workload management, technique optimization, ergonomic adjustment, structured breaks, and recovery may be more appropriate. Previous work has similarly suggested that dividing challenging passages into shorter practice sessions and distributing them throughout the day can support adequate rest and recovery (Chan & Ackermann, 2014).

Third, the substantial between-subject variability observed in many exposures indicates that interventions should not rely solely on generalized recommendations. Individualized approaches that consider technique, posture, instrument setup, and motor strategy may be particularly important for exposures predominantly related to between-subject differences or jointly related to musical characteristics and between-subject differences. This variability may also help explain why assistive devices, such as dynamic supports or ergonomic chin and shoulder rests, have shown at most marginal benefits and, in some cases, adverse effects on muscle activity or perceived exertion (Kok et al., 2019; Mann, Olsen, et al., 2023). These findings highlight the limited effectiveness of interventions that do not account for the complex relationships among musical demands, physical exposure, and individual playing strategy. By distinguishing exposures related primarily to musical characteristics, between-subject differences, or both, the present study provides a basis for developing more targeted and individualized interventions.

Finally, the observed motor patterns, such as coordinated bilateral engagement, redistribution of shoulder muscle activity, and task-specific wrist adjustments, may inform pedagogical approaches in violin teaching. Encouraging efficient movement strategies observed in some experienced musicians may help reduce unnecessary loading and improve long-term musculoskeletal health. More broadly, knowing how musical characteristics affect physical exposures, and how these effects vary across individuals, may help tailor general prevention strategies to the specific repertoire, practice routines, and technical needs of individual violinists.

### 6.5. Strengths and limitations

A key strength of this study is that it systematically examined the influence of three fundamental musical characteristics, i.e., tempo, dynamics, and string, on kinematics and muscle activity in the upper body and upper extremities of professional violinists.

The investigated tasks, i.e. playing scales, incorporated controlled variations in dynamics and string, thus enabling assessment of the influence of each characteristic on biomechanical exposure. However, to maintain a manageable experimental duration, the range of tempi was limited, and not all combinations of tempo, dynamics and string were tested. Specially, the choice of strings was proxied by the scale key, while in practice, there are four strings in combination with multiple main left-hand positions, which was not fully tested with high resolution. As a result, the observed effects, especially the interaction effects between the three characteristics, should only be evaluated to a limited extent. Future studies should investigate a broader range of interactions between musical characteristics to determine whether their effects on physical exposure are independent or combined. Moreover, real musical performance is substantially more complex than scale playing, and the effects observed here may not fully reflect those present when playing an actual repertoire. Future studies should therefore investigate the generalizability of these findings in more ecologically valid musical contexts.

Some participants reported musculoskeletal pain within the preceding 7 days or 12 months. However, all participants were active professional orchestral musicians who did not report pain to be sufficiently severe to require sick leave; thus, they were able to perform as usual. In addition, the prevalence of reported pain in this sample was lower than that typically observed in other populations of musicians (Ioannou et al., 2018; Kenny et al., 2018). Therefore, any influence of pain on performance is likely to have been minimal.

The sample consisted of 12 professional violinists with a mean age of 49 years, which limits the generalizability of the findings. The primary aim of our study was to assess associations between musical characteristics and exposure, rather than to develop fully validated tools for predicting exposure while playing complex pieces of music. Future work should include larger and more diverse samples of string players, and investigate more complex and realistic musical tasks, in order to develop predictive models and evaluate their practical applicability. Also, studies are needed to examine whether informed interventions for violinists, such as alternations between music pieces of different complexity or changed schedules for rehearsal and concerts, can beneficially change patterns of physical exposures and lead to reduced PRMSDs.

## 7. Conclusion

This study demonstrated that three musical characteristics, i.e., tempo, dynamics, and string, were systematically associated with upper-extremity biomechanical exposures during standardized scale playing by professional violinists. The associations differed across musical characteristics and exposure variables. Variance analysis further revealed that some exposures were predominantly related to musical characteristics, others reflected substantial contributions from both musical characteristics and between-subject differences, and others were predominantly related to between-subject differences or residual variability.

Future research should validate these findings in larger samples and more complex musical contexts to determine whether the observed effects are also evident during the performance of complete musical pieces. Estimating physical exposure from musical characteristics may ultimately support exposure prediction directly from sheet music. Moreover, understanding how musical characteristics influence physical exposure may clarify mechanisms contributing to the risk of PRMSDs among violinists and support the development of targeted interventions, including repertoire-based workload management, optimized rehearsal and practice schedules, structured breaks and recovery, and individualized technique-focused strategies.

## 8. Acknowledgements

The authors thank Professor Mikael Forsman for support and consultancy during the early stages of the project, and PhD Ida-Märta Rhén and PhD student Johan Rydgård for their assistance with the illustration of violin playing. The authors also express their sincere gratitude to all violinists participating in this study.

## 11. CRediT author statement

**Xuelong Fan**: Conceptualization, Data curation, Formal analysis, Investigation, Methodology, Software, Validation, Visualization, Writing – original draft, Writing – review & editing. **Svend Erik Mathiassen**: Conceptualization, Funding acquisition, Investigation, Methodology, Writing – review & editing. **Peter J. Johansson**: Conceptualization, Investigation, Resources, Writing – review & editing. **Jennie A. Jackson**: Conceptualization, Investigation, Methodology, Project administration, Resources, Supervision, Writing – review & editing. **Teresia Nyman**: Conceptualization, Funding acquisition, Investigation, Methodology, Project administration, Supervision, Writing – review & editing.

## 12. Declaration of generative AI and AI-assisted technologies in the manuscript preparation process

Statement: During the preparation of this work the author(s) used ChatGPT in order to improve language and readability. After using this tool/service, the author(s) reviewed and edited the content as needed and take(s) full responsibility for the content of the published article.

13.

## Supplemental material

**Table S1.**
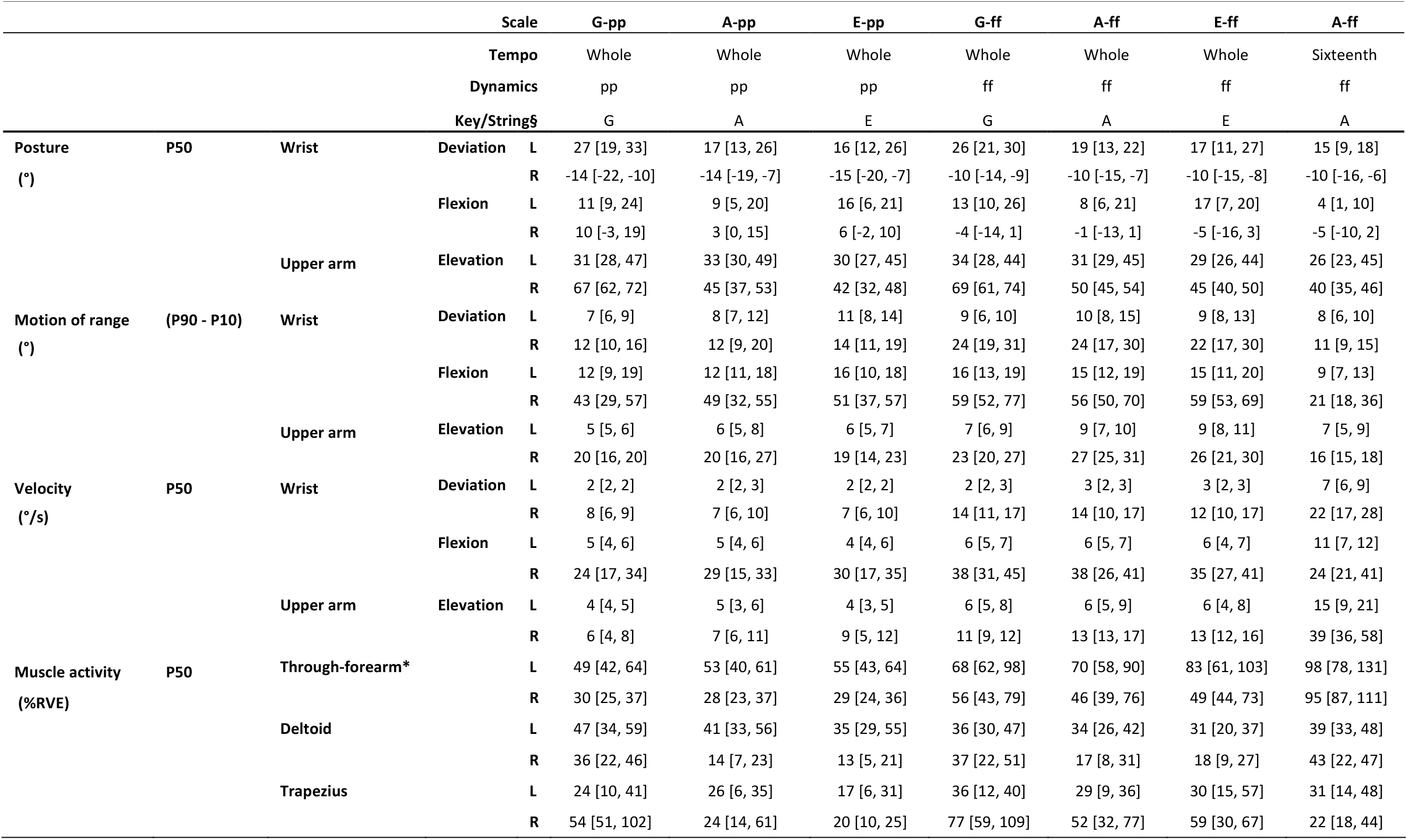
Descriptive results (median [Q1, Q3]) of the exposures in each playing condition. L/R indicates the left or right side; Q1 and Q3 denote the first and third quartiles. * Forearm muscle activity measured by the through-forearm EMG placement. § The scale key was used as a proxy for the strings played during each scale.

**Figure S1.**
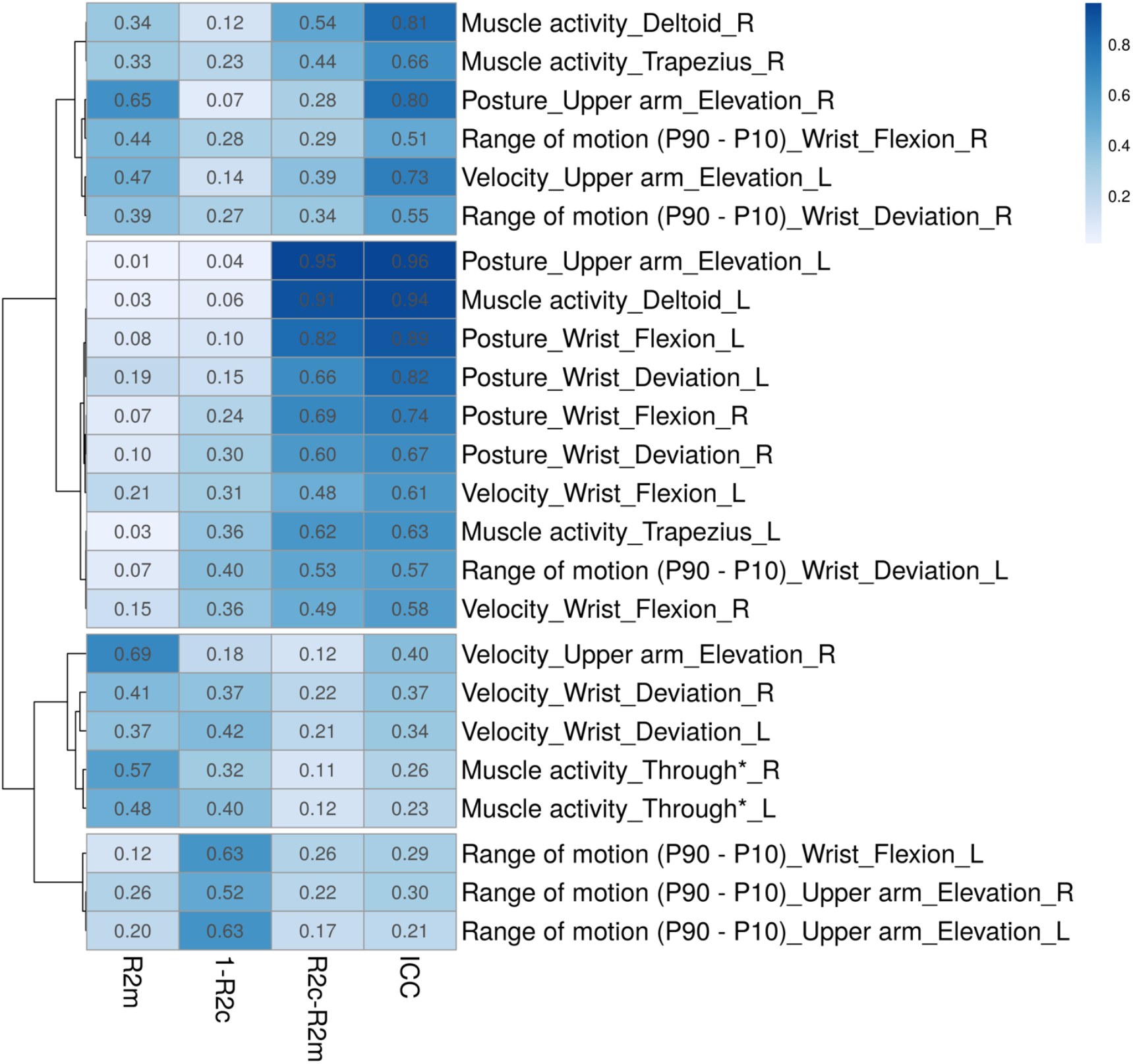
Visualization of variance analysis results of the measured physical exposures. The visualization was generated by ClustVis 2.0 (BIIT, Estonia). A new parameter, 1-R2c, was added to facilitate clustering. No scaling or centering is applied to columns or rows. Rows are clustered using correlation distance and average linkage. The legend shows the magnitude of variance metrics.

